# Head-level lesion-symptom mapping of picture naming in vision-language models

**DOI:** 10.64898/2026.09.15.751760

**Authors:** Samaneh Nemati, Roger D. Newman-Norlund, Saeed Ahmadi, Xiang Guan, Kalil Warren, Yong Yang, Srihari Nelakuditi, Chris Rorden, Leonardo Bonilha, Julius Fridriksson

## Abstract

Researchers in artificial intelligence increasingly intervene on language models to study how their functions are organized, silencing weights and attention components in ways reminiscent of the brain lesions long used to map language in post-stroke aphasia. Under this program of mechanistic interpretability, a common move is to ablate an attention head and read the resulting drop in a behavior as evidence that the head implements it. How much that move can establish about where a behavior is computed remains unclear, because a head whose removal disrupts a behavior is necessary for it but need not be the site where it is computed. We examined this question for the picture naming task, taking the loss of naming (anomia) that defines aphasia as the behavior of interest. Across six vision-language models, spanning three language backbones and a range of parameter scales, we ablated each attention head in turn during single-word picture naming and measured accuracy before and after. The degree of localization varied widely across models. In LLaVA-1.6-Vicuna-13B, a single early head (layer 0, head 20) was necessary: removing it alone reduced naming accuracy from 0.99 to 0.006. The same head was not sufficient, because retaining it while ablating the other 1,599 heads also produced 0% accuracy. Two Mistral-backbone models (LLaVA-Mistral-7B and Idefics2-8B) had no critical head. An early-layer dependence was present in every model but varied in strength, whereas dependence on any single head ranged from dominant to absent. Within Qwen2.5-VL, a dominant head was present in the 7B model but not the 3B model, indicating that this concentration emerged with scale rather than being fixed across a model family, and it was not explained by attention type. These results show that ablating the head whose removal disrupts naming does not establish that the head computes the behavior, that the result generalizes across models, or that the behavior localizes to a head at all.

## 1. Introduction

For more than a century, the study of language has relied on the logic of the lesion. When damage to a brain region disrupts a function, the region is inferred to support that function. This reasoning organized the classical neurology of language, from the deficit-to-region correlations of Broca and Wernicke [1, 2, 3] to their modern statistical form in lesion-symptom mapping [4, 5]. Mechanistic interpretability now applies the same reasoning to artificial neural networks built on the transformer architecture [6]. A candidate component, most often an attention head, is switched off, the change in behavior is measured, and a component whose removal degrades the behavior is taken as evidence that the component implements it. Behaviors have been localized in this way to induction heads [7] and to circuits of heads, most influentially the indirect-object-identification circuit in GPT-2 [8], while factual and lexical associations have instead been attributed to the feed-forward sublayers [9, 10].

This inference has limits that are recognized in both fields. Removing a component and observing that a behavior degrades shows that the component is necessary for the behavior. It does not show that the component is sufficient, that the behavior is computed there rather than routed through it, or that a component identified in one system plays the same role in another. The circuit literature already treats necessity and sufficiency together with redundancy. The GPT-2 identification circuit is a distributed *set* of heads that is jointly necessary and sufficient, and it contains backup heads that remain inactive until the primary heads are ablated, so the individual heads are often not necessary even though the set is sufficient [8]. Related work shows that causal localization need not predict where a behavior can be edited [11], that single ablations are not enough without stronger controls [12, 13], and that large fractions of a transformer’s layers can be removed with little loss [14]. That a component can be necessary without being sufficient, and that an attention head is only one mechanism in a layer, working alongside its feed-forward sublayer, are both familiar, and neither is what we claim. The open question is how far a single behavior localizes at all.

This study measures how much picture naming depends on individual attention heads in vision-language models, and whether that dependence is consistent from one model to the next. In each model we presented a line drawing from a clinical naming test (the Philadelphia Naming Test), asked the model to return the object’s name in a single word, and scored the response as correct or incorrect, before and after ablating attention heads. We addressed three questions across six vision-language models. The first was whether picture naming depends on a small number of specific attention heads. The second was whether a head that is necessary for naming is also sufficient for it, that is, whether that head alone can support naming once the others are removed. The third was whether the organization found in one model holds in the others.

Picture naming suited this purpose for reasons that matter both to machine-learning interpretability and to the study of aphasia. Naming has a well-defined input boundary, because these models encode the image separately and inject it as tokens, so visual information must enter at the earliest layers before anything can be named. Naming is also the behavior at the center of the clinical literature on aphasia, because anomia, the failure to retrieve words, is its most common symptom [15, 16, 17]. Confrontation naming is the standard clinical assay of language after stroke. Our group studies aphasia and its recovery, from the lesion-symptom architecture of the language network [18, 5] to trials of stimulation-based treatment [19], and it is that clinical program that makes naming the dependent measure throughout this series.

This is one of several papers in which our group lesions language models through an aphasia lens. In BLUM, we perturb whole transformer layers and project the resulting error profiles into the lesion space of 410 people with chronic post-stroke aphasia, so that the human brain provides an external reference [20]. Companion papers carry the layer-resolved lesion-symptom-mapping inference for these error maps (PRISM [21]) and the inverse fitting of model perturbations to individual patients’ aphasic error profiles [22].

The present study, LOCUS (Localization Of Computational Units in Subnetworks), asks a narrower question than these. It resolves the dependence to individual attention heads rather than whole layers, it pairs each necessity result with a sufficiency test, and it makes no correspondence claim to the brain, treating degree of localization within the models as the object of study. A separate and fast-growing line of work outside our group lesions language models to reproduce aphasic behavior, either by disabling mixture-of-experts or feed-forward components [23, 24] or by ablating parameters across a text model’s layers [25]. That work targets text-only models and aggregate symptom profiles rather than head-level localization of a single task. Head-level lesion mapping in the model therefore speaks at once to a machine-learning question, what ablation evidence does and does not establish, and to a cognitive-neuroscience question, how naming is organized in systems that are at present among the best computational models of the human language network [26, 27, 28].

No single model answers these three questions, because the answer is the variation between models. We find that the number of necessary heads ranges from one to none, that an early stage of vision intake is present in every model while its collapse onto a single head is not, that this concentration emerges with scale within a model family, and that a critical stage can exist without any critical unit. Read together, these results indicate that ablating the head whose removal breaks a function is not sufficient to establish where the computation lives, that the result will generalize, or that the function localizes to a head at all.

## 2. Results

Every model ran the same arc on 158 curated Philadelphia Naming Test images. We switched off a head by zeroing its query-projection rows and its output-projection columns in the language-model self-attention. Under grouped-query attention the shared key and value projections were left intact, so a single query head was removed cleanly. We never touched the vision encoder, the feed-forward blocks, or the layer norms, and we restored all weights between conditions. Answers were scored by the eight-category word-error classifier of Yang et al. [22], a companion paper from our group, of which this study used only the correct-versus-incorrect distinction. Naming accuracy is the fraction correct, and an accuracy below 0.5 is a collapse. The full procedure is given in Methods.

### 2.1. A single necessary-but-not-sufficient gateway in Vicuna-13B

We first establish the phenomenon in its most extreme form, in LLaVA-1.6-Vicuna-13B, a model with 40 layers, 40 heads per layer, and 1,600 heads in total, whose intact naming accuracy is about 0.99.

Naming localizes to the earliest layers. When we slide a four-layer attention-off window across the network, naming collapses only over layers 0 to 3. Every later window leaves naming near the intact ceiling, as does the same four-layer block placed at the back of the network rather than the front (Fig. 1; Fig. S1, Fig. S2).

**Figure 1:**
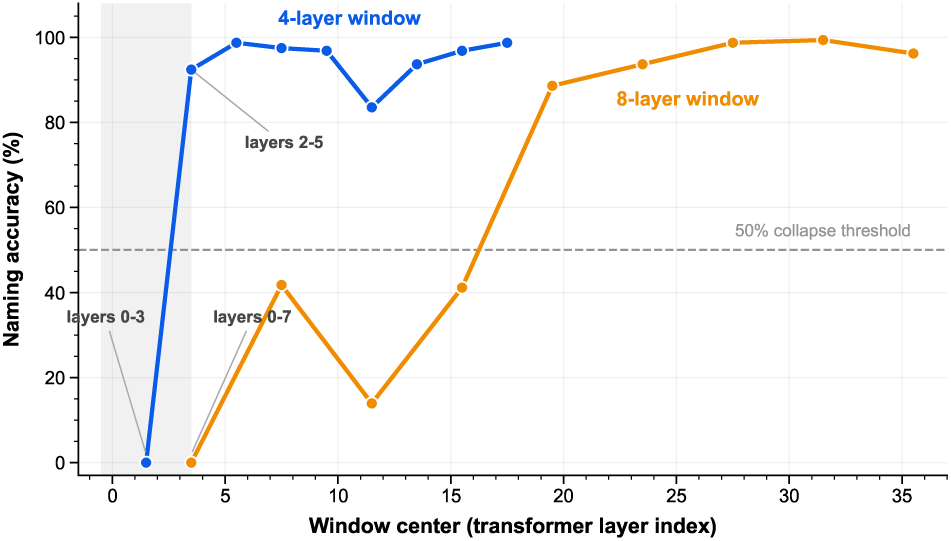
Naming depends selectively on attention in the earliest layers of LLaVA-1.6-Vicuna-13B. Naming accuracy across 158 images is shown after switching off all self-attention within sliding windows of four (blue) or eight (orange) consecutive transformer layers; the horizontal position indicates the center of each window. The shaded region marks the critical layers 0–3. The four-layer lesion collapses naming only when it covers layers 0–3, whereas the overlapping layers 2–5 window preserves near-ceiling accuracy. Larger eight-layer lesions also produce their strongest deficits near the front of the network, while later windows are largely inert.

Inside layers 0 to 3, two observations narrow the effect to a single head. First, when we switch off random subsets of the 160 heads in these layers, it is *which* heads are removed, not how many, that decides whether naming collapses. At every intermediate count the same number of ablated heads produces both intact naming and total collapse, across 30 random draws per count (Fig. 2). Second, one head, layer 0 head 20 (L0h20), is present in 85% of the collapsing draws and in only 3% of the intact ones, and it stands alone on a per-head importance map while every other head in the region is inert (Fig. 3). Removing L0h20 on its own drops naming to 0.006, its strongest regional neighbor L0h3 has no effect (0.975), and of all 40 first-layer heads only head 20 behaves this way (Fig. 4A). No other four-layer band in the network is fragile at any dose. When we run the identical 30-seed sweep on every band from layers 4 to 7 through 36 to 39, not one collapses, whereas layers 0 to 3 collapse 54 times in 180 draws (Fig. 4B). The failure is silent, because the lesioned model returns an empty response rather than a wrong name, which is the signature of cutting an input stage off from its supply.

**Figure 2:**
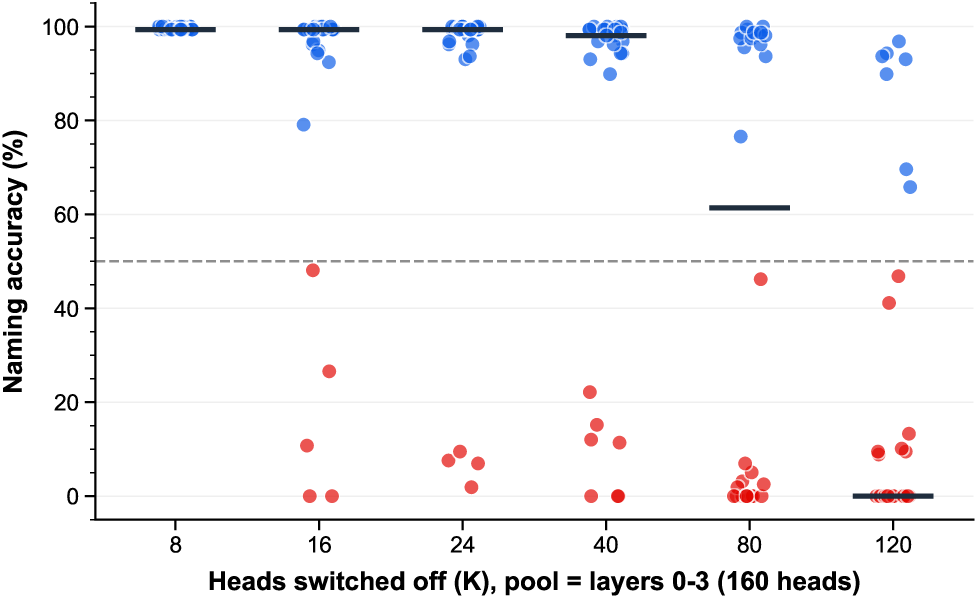
Naming outcome depends on head identity rather than lesion size in LLaVA-1.6-Vicuna-13B. Each point shows naming accuracy across 158 images after ablating a random subset of *K* heads from layers 0–3 (160 heads total; 30 subsets per *K*). Blue points indicate non-collapsing trials and red points indicate collapse (accuracy below 50%); the dashed line marks this threshold, and black bars show medians. For the same *K*, accuracy ranges from near ceiling to complete collapse, demonstrating that the outcome depends on which heads are removed, not simply how many.

**Figure 3:**
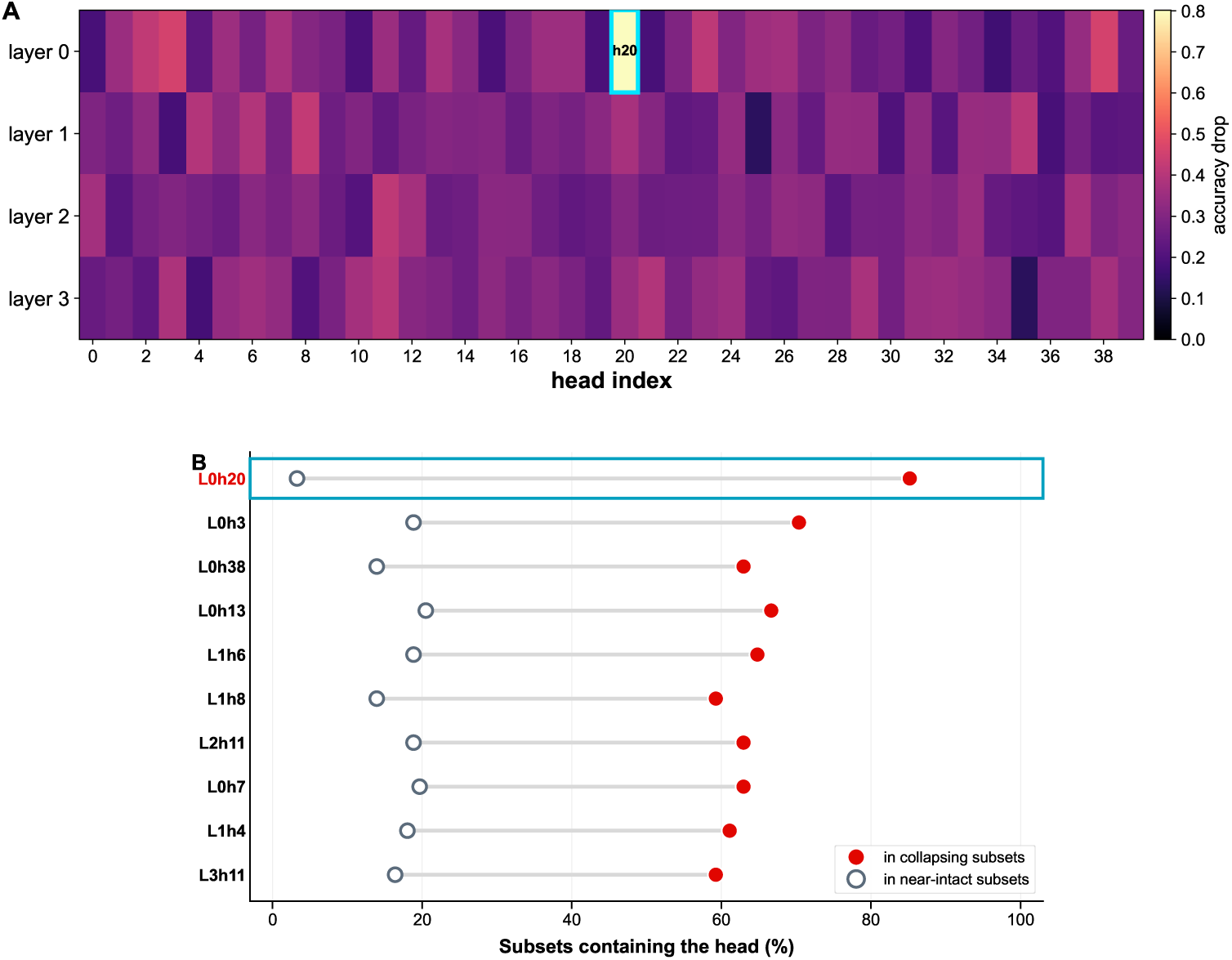
Random-subset ablations identify L0h20 as the dominant head within the critical early block of LLaVA-1.6-Vicuna-13B. Results summarize 180 random-subset ablations across the 160 heads in layers 0–3. (A) Per-head importance, defined as the mean accuracy loss associated with including each head in the ablated subset; L0h20 (boxed) produces a markedly larger loss than any other head. (B) For the ten most enriched heads, the fraction of collapsing subsets (filled, accuracy below 50%) and of near-intact subsets (open, accuracy at least 85%) that contain each head. L0h20 appears in 85% of collapsing subsets but in only 3% of near-intact ones, a separation at both ends that no other head shows.

**Figure 4:**
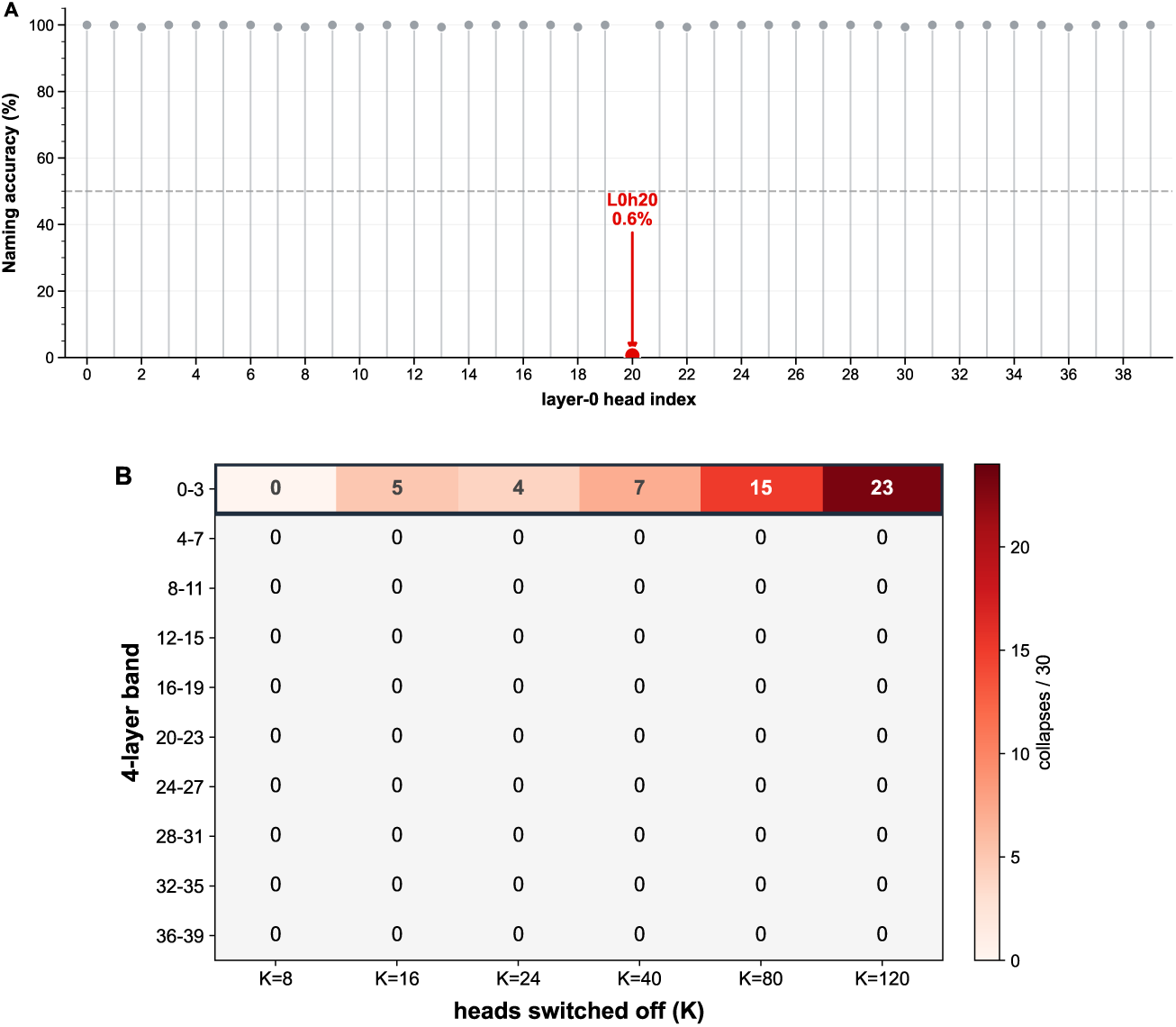
The single-head gateway is sharp and confined to the earliest layers in LLaVA-1.6-Vicuna-13B. (A) Naming accuracy across 158 images after ablating each of the 40 layer-0 heads individually. Only L0h20 causes collapse (0.6% accuracy); removal of any other head preserves near-ceiling performance (99–100%). The dashed line marks the 50% collapse threshold. (B) Number of collapsing trials among 30 random subsets of *K* ablated heads within each nonoverlapping four-layer band. Collapses occur only in layers 0–3 and become more frequent as *K* increases; no later band collapses at any tested lesion size.

The decisive test is sufficiency. When we keep only L0h20 and switch off the other 1,599 heads, naming also falls to 0% (Table 1). The one head that is individually required cannot, on its own, name at all. Read together, the two lesions identify L0h20 as an indispensable early gateway, not as a unit that performs naming. The same collapse appears under three mechanistically distinct interventions. An inference-time forward-hook knockout that leaves the weights intact reproduces it under both zero-ablation and mean-ablation, all to 0.000. Because mean-ablation removes the head’s image-specific information while preserving its average contribution, this shows that the head carries image-specific information rather than a constant bias (Table 1). Scaling the head down rather than off yields a graded dose-response in accuracy. Naming falls to zero under a heavy lesion and returns to the intact ceiling by about 60 to 65% of full strength, a curve that is stable across decoding seeds (Fig. S3). The head’s setting, and not merely its presence, controls how much naming survives.

**Table 1:** L0h20 is necessary but not sufficient, and intervention-invariant (Vicuna-13B, 158 images).

| Lesion | Naming accuracy | Reads as |
| --- | --- | --- |
| Remove L0h20 (weight) | 0.006 | necessary |
| Remove L0h20 (activation, zero) | 0.000 | necessary, weights intact |
| Remove L0h20 (activation, mean) | 0.000 | carries image-specific information |
| Keep only L0h20, remove other 1,599 | 0.000 | not sufficient |
| Remove L0h3 (neighbor control) | 0.975 | specific |

### 2.2. Head-level localization ranges from a single critical head to none across six models

We then ran the identical arc on five further models, spanning two more lineages and a range of sizes, and asked the same counting question of each. The answer is a spectrum (Table 2, Fig. 5). At one extreme, Vicuna-13B reduces to a single clean head. Qwen2.5-VL-7B, from a different lineage, also concentrates the early dependence into a dominant head (L0h22, 0.006 when removed alone), but it has three weaker companion heads in the same layer (0.057, 0.222, 0.386), so the head is dominant rather than cleanly single. Vicuna-7B has a single head at a milder severity (L0h3, 0.158). At the other extreme, LLaVA-Mistral-7B and Idefics2-8B have no critical head at all, because every single-head ablation leaves naming at or near the intact baseline (at or above 0.949 and 0.930 respectively). The single-head phenomenon is therefore real but not universal, and even where it appears it is not always clean.

**Figure 5:**
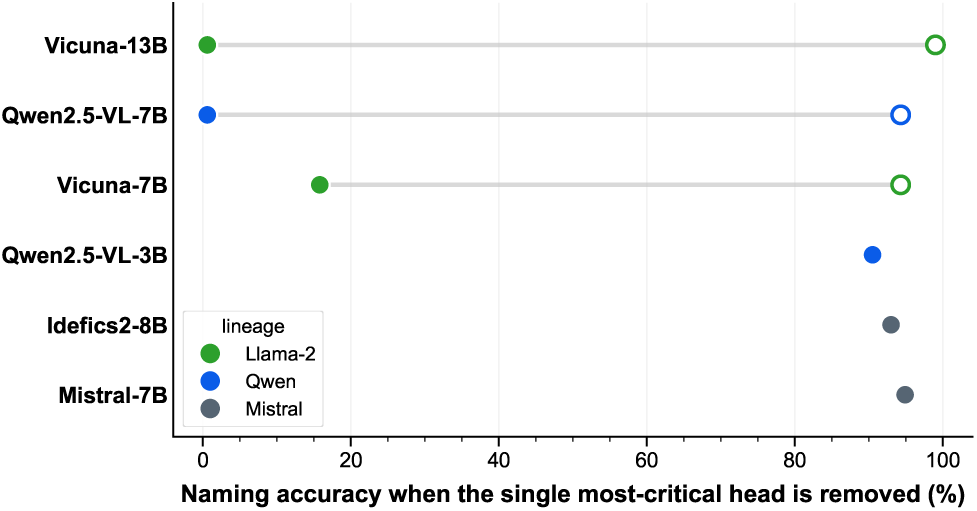
The degree of head-level localization varies across models. Filled circles show naming accuracy across 158 images after ablating the single head that produces the largest accuracy loss in each model; open circles show intact-model accuracy, and horizontal lines connect the paired values. Models are ordered from strongest to weakest single-head dependence. A single-head ablation nearly abolishes naming in Vicuna-13B and Qwen2.5-VL-7B, partially impairs it in Vicuna-7B, and has no measurable effect in Qwen2.5-VL-3B, Idefics2-8B, or Mistral-7B, where the filled and open circles overlap. Colors indicate language-model lineage.

**Table 2:** Six models on the same arc. *Intact*, naming accuracy of the undamaged model. *Early block critical*, whether the earliest layers are jointly necessary. *Critical single head*, the head whose removal alone most reduces naming, with the resulting accuracy in parentheses; “none” means no single head is critical.

| Model | Lineage | Intact | Early block | Critical single head |
| --- | --- | --- | --- | --- |
| Vicuna-13B | Llama-2 | $\sim 0.99$ | yes | L0h20 (0.006) |
| Qwen2.5-VL-7B | Qwen | 0.943 | yes | L0h22 (0.006), +3 weaker |
| Vicuna-7B | Llama-2 | 0.943 | yes | L0h3 (0.158) |
| Qwen2.5-VL-3B | Qwen | 0.905 | yes | none ($\geq 0.905$) |
| Idefics2-8B | Mistral | 0.930 | weak | none (0.930) |
| Mistral-7B | Mistral | 0.949 | no | none ($\geq 0.949$) |

### 2.3. Early-layer dependence is common, but concentration in a single head is not

What is common to all six models is an early vision-intake bottleneck: naming depends on the earliest layers taken as a block. In every model, ablating a large enough early window drives naming to zero, for example layers 0 to 3 in Qwen-3B and layers 0 to 7 in Qwen-7B and Idefics2, while later windows are largely inert. What differs is whether that block reduces to a single head. Qwen2.5-VL-3B is the decisive case, because it separates the two. Its first four layers are jointly necessary, since ablating layers 0 to 3 gives 0.000, and yet switching off each of the 64 heads in those layers on its own leaves naming at the intact 0.905 in every case. A 30-seed random-subset analysis shows that the dependence is carried diffusely, with the probability of collapse rising as more early heads are removed (0, 5, 12, 27, and 30 collapses out of 30 at 8, 16, 24, 32, and 48 heads) and no single head enriched. A critical processing stage need not contain a critical unit.

### 2.4. Single-head concentration differs with scale within Qwen2.5-VL

Because Qwen provides two sizes of one architecture, it lets us separate scale from lineage. The 3B model has the early bottleneck but no critical head, while the 7B model has a dominant critical head in the same early layer. With the same recipe and the same vision-injection scheme, and only the size changed, the concentration of the bottleneck onto a single head is present in the larger model and absent in the smaller one, so it emerges with scale rather than being a fixed property of the family (Fig. 5, Table 2). Attention type does not explain the pattern, because Qwen and Idefics2 both use grouped-query attention and yet sit at opposite ends of the spectrum, so grouped-query attention is not what removes the gateway. The factor that tracks concentration is some combination of scale and language backbone, and the step from Qwen-3B to Qwen-7B is only the first rung of the size ladder that would separate them.

## 3. Discussion

### 3.1. Head-level localization varies across models for the same task

The organizing result is that, for a fixed task, how localized the computation is varies from one model to the next, from a single head carrying the entire dependency to no single head mattering at all. This changes what a head-ablation result can mean. In the identification-circuit tradition, one finds within a single model a distributed set of heads that is necessary and sufficient, with redundancy and backup heads [7, 8], and that program answers the question of what the circuit is in that model. A neuroscience-styled variant localizes a distributed set of language-selective units in text language models whose ablation causes large deficits [29]. Our question is different and comparative: how localized is this function, and does the answer generalize? The answer is that it does not. The same probe yields a single clean head, a dominant head with companions, a milder single head, a critical block with no critical head, or nothing at all, depending on the model. Treating degree of localization as a measured quantity rather than an assumption is the contribution here, and the six models are the measurement, not a robustness afterthought.

### 3.2. Why L0h20 is a gateway, not a naming module

The two lesions that bound the Vicuna-13B result, necessary on removal and insufficient in isolation, are what license the gateway reading, and two features of the data favor it over a reading in which the head is a naming unit. The failure is silent rather than error-laden, which is the signature of starving an input stage rather than corrupting a computation, and it fits naming having a literal input boundary where image tokens enter the earliest layers. The graded lesion produces a smooth accuracy dose-response rather than an abrupt switch, which is the behavior of a channel whose throughput sets how much visual evidence reaches an intact downstream namer, not of a partially damaged namer. A single dial that moves naming from silence to ceiling is also why this head can serve as a severity control in the companion digital-twin work [22].

### 3.3. Why a robust single-head effect can still mislead

The gateway is confirmed by three mechanistically distinct interventions, weight zeroing, activation zero-ablation, and activation mean-ablation, and by a seed-stable graded dose-response, so it is unlikely to be an artifact of any one intervention. Taken together with the cross-model spectrum, the results point to three distinct ways in which reading a single head-ablation as a mechanism can fail. First, the head can be necessary without being where the computation happens, as when it is an input relay, which is what the keep-only test shows for L0h20. Second, the finding can fail to generalize, because the same task localizes to a single head in some models and to none in others, so a circuit identified in one model is not a fact about the architecture, in line with earlier evidence that mechanistic roles need not transfer. Third, the function may not reduce to any head at all, as in Qwen-3B, which has a critical early stage but no critical head. Each of these is a separate caution, and together they say that necessity by ablation is the beginning of an argument about mechanism, not its conclusion. The corrective is inexpensive, and we recommend it as routine: pair every necessity claim with a sufficiency probe (keep-only), a graded intervention, and, where possible, a second model, before describing a component as performing a function [12, 13, 11].

### 3.4. Possible origins of single-head concentration: scale, lineage, and architecture

Our clearest positive signal is that single-head concentration may increase with scale within the Qwen family, although this rests on a single 3B-to-7B step and we are cautious about reading a single cause into it. Concentration appears in two lineages, Llama-2 and Qwen, and is absent in a third, Mistral, across two models that differ in connector; within Qwen, the 7B has a dominant head while the 3B does not. Scale and backbone are both implicated, and the present data cannot fully separate them, since scale, backbone, connector, and training history are partly confounded across these six models. That the early bottleneck itself is invariant across all six models, while only its concentration varies, suggests the bottleneck is a structural consequence of injecting vision as tokens at the front of the network, a hypothesis a cross-attention-injection model would test.

### 3.5. Where model lesions and human aphasia converge and diverge

In humans, picture naming is not gated by a single indispensable bottleneck whose loss silences all naming. Damage instead produces graded, category-patterned anomia across a distributed ventral system [17, 18, 5]. The models that do concentrate naming into one early head do so because their architecture funnels vision into language at a fixed location, which the brain does not. Naming this divergence is more useful than papering over it. The model-as-brain correspondence is well supported at the representational level [26, 27, 28], but it breaks at the input interface, and it breaks to a degree that itself depends on the model, which limits how far any single model’s lesion results can be carried back to clinical inference. This is where the present work and BLUM [20] meet from opposite directions. BLUM shows that model error profiles, at the level of transformer layers, project onto human lesion patterns above chance, with semantic errors tracking ventral-stream damage and phonemic errors tracking dorsal-stream damage. LOCUS looks one level finer and finds the single early head that gates all naming, which has no counterpart in a real brain. The two are complementary, one establishing the correspondence at layer resolution and the other showing where it breaks at head resolution.

The task, stimuli, and scoring used here are the clinical instruments of anomia: single-word naming of Philadelphia Naming Test pictures, scored with the taxonomy that aphasia assessment uses. That shared measurement is what lets a model result speak to aphasia at all, and it gives the two main findings a clinical reading. The graded gateway is a controllable severity axis for naming, which is exactly the ingredient a patient-specific digital twin of anomia needs in order to match an individual’s impairment. The range from one head to none is a caution in the other direction, because how naming is organized, or how it fails, read from one model need not hold in another, and still less in a patient. LOCUS models no patient. It maps what naming does under lesion in the models the companion patient work is built on, and it marks where that behavior stops resembling a brain.

## 4. Conclusion

We asked how strongly picture naming depends on individual attention heads and whether that dependence generalizes across vision-language models. It does not. Across six models, the effect of removing a single head ranged from near-total loss of naming to no measurable impairment. In LLaVA-1.6-Vicuna-13B, removing L0h20 nearly abolished naming, yet retaining that head while removing the other 1,599 heads could not support naming at all. L0h20 is therefore a necessary early gateway, not the location where naming is performed.

Dependence on the earliest layers was more consistent across models than dependence on any individual head, although its strength varied. Whether that early dependence became concentrated in a single head differed across model families and sizes. Within Qwen2.5-VL, a dominant critical head appeared in the 7B but not the 3B model, suggesting that scale may contribute to this concentration, although additional model sizes are needed to establish that relationship.

Removing a component and observing behavioral failure establishes necessity, but not sufficiency, computational location, or generality across models. Before describing an attention head as performing a function, the ablation result should be paired with a sufficiency test, examined with a graded intervention, and tested across additional models. Attention alone is not enough to explain where a behavior is computed.

## 5. Methods

### 5.1. Models

We used six open-weight vision-language models, all of which inject vision as tokens into a transformer language model: LLaVA-1.6-Vicuna-13B and –7B [30, 31], LLaVA-1.6-Mistral-7B, Qwen2.5-VL-3B and –7B [32], and Idefics2-8B [33]. The two Vicuna models are Llama-2 fine-tunes wrapped in the LLaVA late-fusion design, LLaVA-Mistral-7B and Idefics2-8B are built on a Mistral-7B backbone, and Qwen2.5-VL is natively multimodal. We loaded each model in half precision, and read its layer count, head count, key/value-head count, and head dimension from its configuration. We ran the Vicuna and Mistral LLaVA models through the shared LOCUS pipeline. For Qwen2.5-VL and Idefics2 we used self-contained per-model ports that reproduced the pipeline through each model’s own interface. For Idefics2 we pinned the lesion to the text decoder, because the perceiver connector has its own self-attention at layer indices that overlap the text layers, and early-layer targeting must not touch it. Generation was greedy, with a one-word naming prompt in each model’s chat template.

### 5.2. Task and scoring

Each model named each Philadelphia Naming Test image in a single word. We scored results on the 158-item curated subset used throughout the LOCUS series. Each answer was labelled by the eight-category word-error classifier of Yang et al. [22], and this study used only the correct-versus-incorrect distinction, so naming accuracy is the fraction labelled correct and a collapse is an accuracy below 0.5. We decoupled generation from scoring, writing answers to CSV and scoring them afterward.

### 5.3. Head interventions

We switched off a single query head by zeroing its rows in the query projection and its columns in the output projection of the language-model self-attention. Under grouped-query attention the key and value projections are shared across a group of query heads, so we left them intact and zeroed only the query and output slices, which removed one query head cleanly without silencing its group. A graded lesion multiplied those same slices by a factor between 0 and 1, where 0 is full removal and 1 is intact. We cloned the targeted weights before a condition and restored them afterward, so conditions were independent and the model was never retrained. We never modified the vision encoder, the feed-forward blocks, or the layer norms.

### 5.4. Experimental conditions and ablation sweeps

We ran the same conditions for each model. After the intact baseline, we zeroed all attention in all layers to confirm that attention is necessary for naming (0% in Vicuna-13B). A centered dose sweep varied the size of an attention-off block from 1 to 30 layers, refined between 11 and 14. For localization we placed a fixed 10– and 13-layer block at the front, middle, and back, then stepped an 8-layer window across all layers and a 4-layer window across the early region in steps of two, which separated position from dose. At head resolution, within the critical early block (layers 0 to 3 in Vicuna-13B, 160 heads), we switched off random subsets of 8, 16, 24, 40, 80, and 120 heads with 30 draws each (180 conditions), from which we computed per-head importance and collapse-enrichment, followed by a single-head sweep of every head in the block. For Vicuna-13B we additionally ran the keep-only sufficiency lesion (keep the candidate head, zero the other 1,599), an inference-time forward-hook knockout of L0h20 under zero– and mean-ablation (with the mean estimated over 40 images), and a graded strength sweep across six decoding seeds. As a control, we repeated the full random-subset grid on every other 4-layer band from layers 4 to 7 through 36 to 39.

### 5.5. Compute and code

Runs used single NVIDIA L40S GPUs at the University of South Carolina. The per-model runners and the cross-model report are in the LOCUS repository (https://github.com/samnemati/locus), and the full per-condition outputs and figure code are released with the paper.

## 6. Limitations

These are single runs per condition, on one naming task, scored by one classifier. The single-head sweeps establish that no individual head is necessary in the distributed models, but they do not rule out a small fixed set in Idefics2, though the Qwen-3B subset analysis argues against a small set there. The scale claim rests on a single 3B-to-7B step within Qwen. The Mistral-lineage negative rests on two models that share a backbone but differ in their connector. Two axes were deliberately held roughly constant, and both are natural next tests. The first is task, since all of our results are for naming, and whether the same gateway governs other model outputs is the single most valuable follow-up, because a genuine input gateway should gate them all. The second is the vision-injection scheme, since all six models inject vision as tokens, and a cross-attention model such as Llama-3.2-Vision would test whether the early bottleneck depends on the injection design. Finally, the method needs open weights and enough compute to run the model, so the largest closed or frontier systems cannot be tested this way.

## Data and code availability

Code, per-model runners, the cross-model report, figures, and all per-condition scored outputs are in the LOCUS repository (https://github.com/samnemati/locus). The naming images are the Philadelphia Naming Test set and are not redistributed; the classifier is that of Yang et al. [22].

## Funding

Supported by the National Institute on Deafness and Other Communication Disorders (NIDCD), including the Center for the Study of Aphasia Recovery (C-STAR). The funders had no role in study design, analysis, or the decision to publish.

## Competing interests

R.D.N.-N. and J.F. have a financial interest in ALLT.AI, LLC. The remaining authors declare no competing interests.

## Author contributions

Conceptualization: S. Nemati, R.D.N.-N., J.F. Methodology: S. Nemati, R.D.N.-N., S. Nelakuditi, C.R. Software: S. Nemati, S.A., X.G., Y.Y. Formal analysis: S. Nemati, R.D.N.-N. Investigation: S. Nemati, R.D.N.-N. Visualization: S. Nemati, R.D.N.-N. Writing, original draft: S. Nemati, R.D.N.-N. Writing, review and editing: all authors. Supervision: J.F., C.R., S. Nelakuditi. Funding acquisition: J.F., C.R.

## Supplementary figures

**Figure S1:**
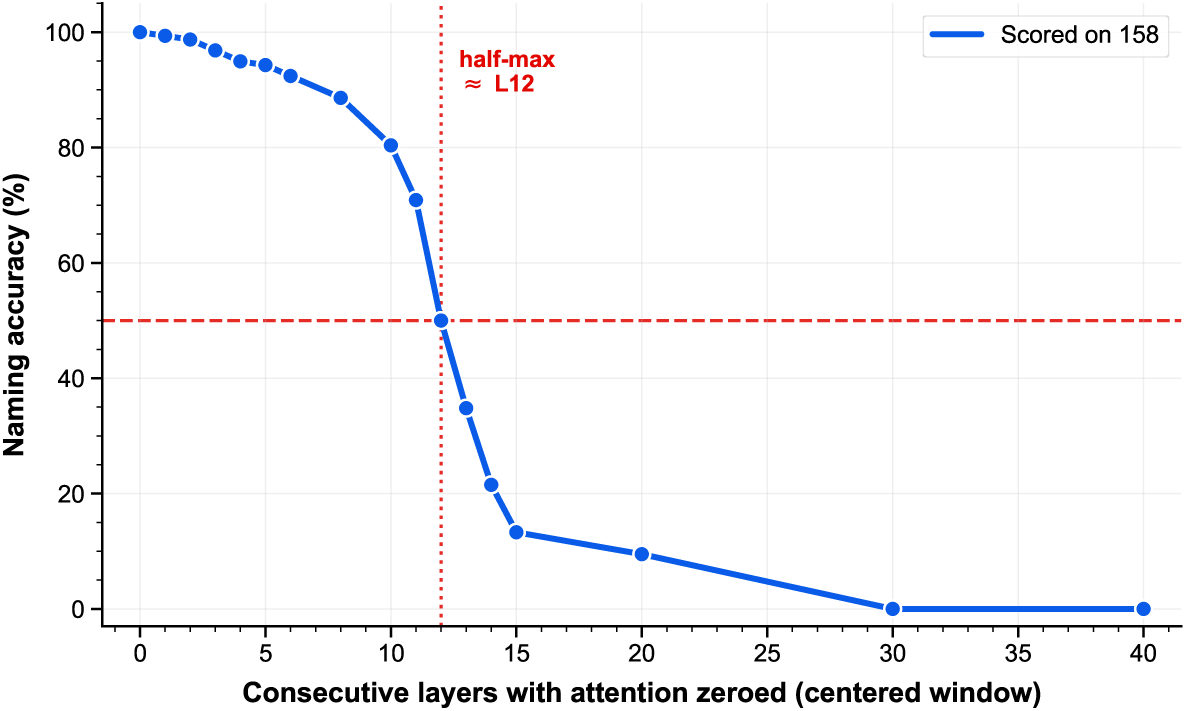
Naming exhibits a dose-dependent decline as progressively larger attention blocks are removed in LLaVA-1.6-Vicuna-13B. Naming accuracy across 158 images is shown after switching off attention in a centered block of *L* consecutive layers. Accuracy declines gradually for small lesions and then sharply, crossing the 50% threshold at approximately *L* = 12 of 40 layers before approaching zero for larger lesions. Dashed lines mark 50% accuracy and the corresponding lesion size.

**Figure S2:**
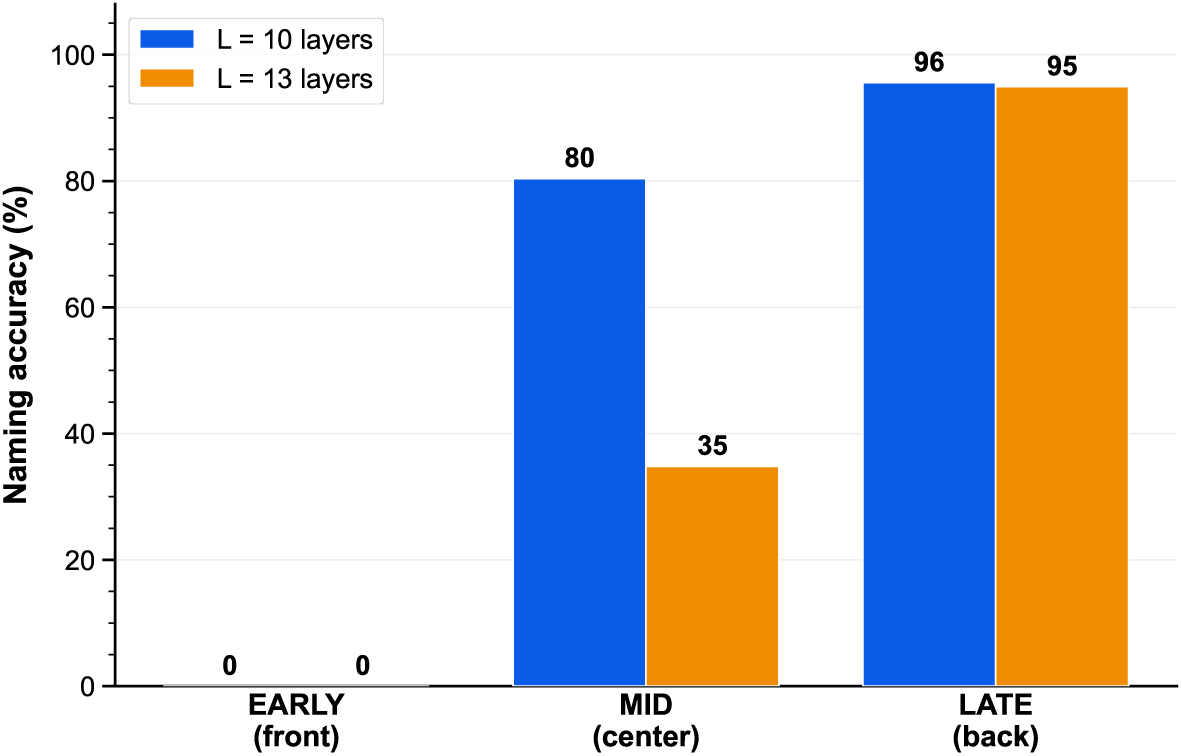
Lesion position, rather than lesion size alone, determines naming failure in LLaVA-1.6-Vicuna-13B. Naming accuracy across 158 images is shown after switching off attention in blocks of 10 or 13 consecutive layers placed at the front, center, or back of the network. Both lesion sizes abolish naming when placed at the front, produce size-dependent impairment at the center, and leave naming near ceiling when placed at the back. Thus, equally sized attention lesions have markedly different effects depending on their location.

**Figure S3:**
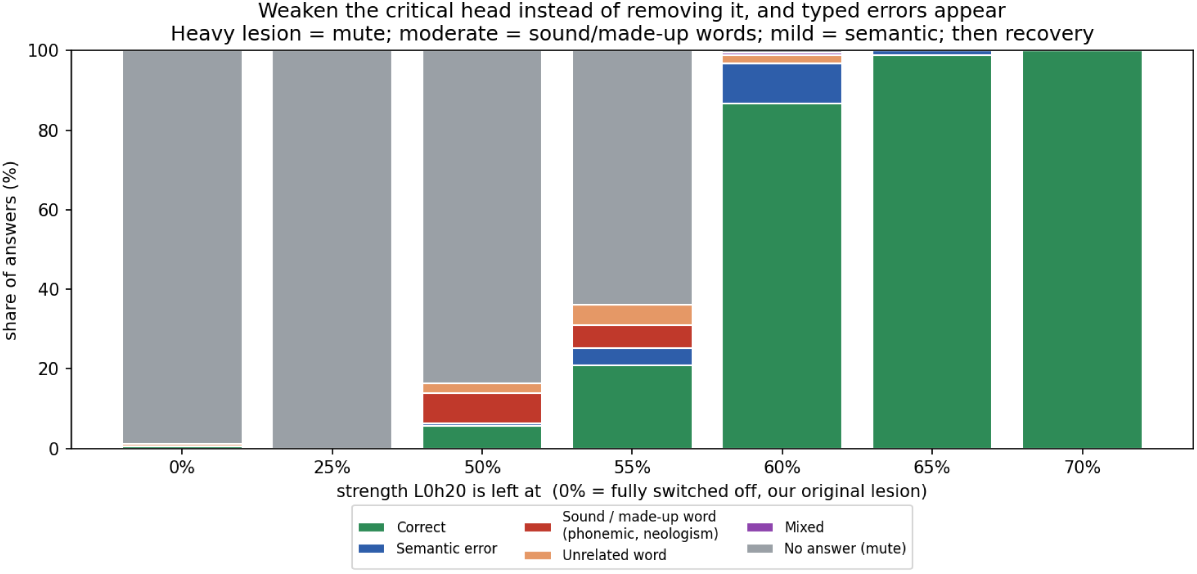
Graded weakening of L0h20 produces an ordered transition from no response to naming errors and then recovery in LLaVA-1.6-Vicuna-13B. Stacked bars show the proportion of responses in each naming-outcome category across 158 Philadelphia Naming Test images as L0h20 is retained at increasing fractions of its intact strength (0% = complete ablation). No response dominates at 0–25% strength, error types (phonemic, made-up-word, semantic, mixed) emerge at intermediate strengths, and correct naming becomes dominant by 60% and near ceiling by 65–70%.

